# Deep learning with EEG spectrograms in rapid eye movement behavior disorder

**DOI:** 10.1101/240267

**Authors:** Giulio Ruffini, David Ibañez, Marta Castellano, Laura Dubreuil, Jean-François Gagnon, Jacques Montplaisir, Aureli Soria-Frisch

**Author notes:** http://neuroelectrics.com, http://starlab-int.com (Giulio Ruffini).

## Abstract

REM Behavior Disorder (RBD) is now recognized as the prodromal stage of *α*-synucleinopathies such as Parkinson’s disease (PD). In this paper, we describe deep learning models for diagnosis/prognosis derived from a few minutes of eyes-closed resting electroencephalography data (EEG) collected from idiopathic RBD patients (n=121) and healthy controls (HC, n=91). A few years after the EEG acquisition (4 ± 2 years), a subset of the RBD patients eventually developed either PD (n=14) or Dementia with Lewy bodies (DLB, n=13), while the rest remained idiopathic. We describe first a simple convolutional neural network (DCNN) with a five-layer architecture combining filtering and pooling, which we train using stacked multi-channel EEG spectrograms. We treat the data as in audio or image classification problems where deep networks have proven highly successful by exploiting compositional and translationally invariant features in the data. For comparison, we study an even simpler deep recurrent neural network using three stacked Long Short Term Memory network (LSTM) cells or gated-recurrent unit (GRU) cells—with very similar results. The performance of these networks typically reaches 80% (±1%) classification accuracy in the balanced HC vs. PD-outcome classification problem. In particular, using data from a single EEG channel we obtain an area under the curve (AUC) of 87% (±1%) while avoiding spectral feature selection. The trained classifier can also be used to generate synthetic spectrograms to study what spectrogram features are relevant for classification, pointing to the presence of theta band bursts and a decrease of power in the alpha band in future PD or DLB patients compared to HCs. We conclude that deep networks may provide a key tool for the analysis of EEG dynamics even from relatively small datasets and enable the delivery of new biomarkers.

## 1 Introduction

RBD is a parasomnia characterized by intense dreams with during REM sleep without muscle atonia [10], i.e., with vocalizations and body movements. Idiopathic RBD occurs in the absence of any neurological disease or other identified cause, is male-predominant and its clinical course is generally chronic progressive [6]. Several longitudinal studies conducted in sleep centers have shown that most patients diagnosed with the idiopathic form of RBD will eventually be diagnosed with a neurological disorder such as Parkinson disease (PD) or dementia with Lewy bodies (DLB) [16, 6, 10, 9]. In essence, idiopathic RBD has been suggested as a prodromal stage of *α*-synucleinopathies (PD, DLB and less frequently multiple system atrophy (MSA) [10, 9]).

RBD has an estimated prevalence of 15–60% in PD and has been proposed to define a subtype of PD with relatively poor prognosis, reflecting a brainstem-dominant route of pathology progression (see [12] and references therein) with a higher risk for dementia or hallucinations. PD with RBD is characterized by more profound and extensive pathology—not limited to the brainstem—, with higher synuclein deposition in both cortical and sub-cortical regions.

Electroencephalographic (EEG) and magnetoencephalographic (MEG) signals contain rich information associated with functional processes in the brain. To a large extent, progress in their analysis has been driven by the study of spectral features in electrode space, which has indeed proven useful to study the human brain in both health and disease. For example, the “slowing down” of EEG is known to characterize neurodegenerative diseases [5, 17, 20]. It is worth mentioning that the selection of disease characterizing features from spectral analysis is mostly done manually after an extensive search in the frequency-channel domain.

However, neuronal activity exhibits non-linear dynamics and non-stationarity across temporal scales that cannot be studied properly using classical approaches. Tools capable of capturing the rich spatiotemporal hierarchical structures hidden in these signals are needed. In [20], for example, algorithmic complexity metrics of EEG spectrograms were used to derive information from the dynamics of EEG signals in RBD patients, with good results, indicating that such metrics may be useful per se for classification or scoring. However, ideally we would like to use methods where the relevant features are found directly by the algorithms.

Deep learning algorithms are designed for the task of exploiting compositional structure in data [15]. In past work, for example, deep feed-forward autoencoders have been used for the analysis of EEG data to address the issue of feature selection, with promising results [13]. Interestingly, deep learning techniques, in particular, and artificial neural networks in general are themselves bio-inspired in the brain—the same biological system generating the electric signals we aim to decode. This suggests they may be well suited for the task.

Deep recurrent neural networks (RNNs), are known to be potentially Turing complete [18], but general RNN architectures are notoriously difficult to train [7]. In this regard, it is worth mentioning that “reservoir” based RNN training approaches are evolving [14]. In related work, a particular class of RNNs called Echo State Networks (ESNs) that combine the power of RNNs for classification of temporal patterns and ease of training [19] was used with good results. The main idea behind ESNs and other “reservoir computation” approaches is to use semi-randomly connected, large, fixed recurrent neural networks where each node/neuron in the reservoir is activated in a non-linear fashion. The interior nodes with random weights constitute what is called the “dynamic reservoir” of the network. The dynamics of the reservoir provides a feature representation map of the input signals into a much larger dimensional space (in a sense much like a kernel method). Using such an ESN, an accuracy of 85% in a binary, class-balanced classification problem (healthy controls versus PD patients) was obtained using a relatively small dataset [19]. The main limitations of this approach, in our view, are the computational cost of developing the reservoir dynamics of large random networks and the associated need for feature selection (e.g., which subset of frequency bands and channels to use as inputs to simplify the computational burden).

In this paper we use a similar but simpler strategy as the one presented in [28], using Convolutional Networks with EEG signals, i.e. multi-channel time series. In contrast to [28], we reduce the number of hidden layers from 16 to 4, use a simpler approach for the generation of spectrograms, and do not use transfer learning from a network trained on a visual recognition task. Indeed, we believe such a pre-training would initialize the filtering weights to detect object-like features not present in spectrograms. The proposed methodology outperforms several shallow methods used for comparison as presented in the results section. Lastly, we propose the utilization of deep-learning visualization techniques for the interpretation of results. This is important for the use and acceptance of such techniques in the clinical domain, where black-box approaches have been extensively criticized,

### 1.1 Deep learning in the spectrogram representation

Here we explore first a deep learning approach inspired by recent successes in image classification using deep convolutional neural networks (DCNNs), designed to exploit invariances and capture compositional features in the data (see e.g., [7, 15, 18]). These systems have been largely developed to deal with image data, i.e., 2D arrays, possibly from different channels, or audio data (as in [27]), and, more recently, with EEG data as well [25, 28]. Thus, inputs to such networks are data cubes (multichannel stacked images). In the same vein, we aimed to work here with the spectrograms of EEG channel data, i.e., 2D time-frequency maps. Such representations represent spectral dynamics as essentially images with the equivalent of image depth provided by multiple available EEG channels (or, e.g., current source density maps or cortically mapped quantities from different spatial locations). Using such representation, we avoid the need to select frequency bands or channels in the process of feature selection. This approach essentially treats EEG channel data as an audio file, and our approach mimics similar uses of deep networks in that domain.

RNNs can also be used to classify images, e.g., using image pixel rows as time series. This is particularly appropriate in our case, given the good performance we obtained using ESNs on temporal spectral data. We study here also the use of stacked architectures of long-short term memory network (LSTM) or gated-recurrent unit (GRU) cells, which have shown good representational power and can be trained using backpropagation [8, 7, 3].

Our general assumption is that some relevant aspects in EEG data from our datasets are contained in compositional features embedded in the time-frequency representation. This assumption is not unique to our particular classification domain, but should hold of EEG in general. In particular, we expect that deep networks may be able to efficiently learn to identify features in the time-frequency domain associated to bursting events across frequency bands that may help separate classes, as in “bump analysis" [4]. Bursting events are hypothesized to be representative of transient synchrony of neural populations, which are known to be affected in neurodegenerative diseases such as Parkinson’s or Alzheimer’s disease [26].

Finally, we note that we make no attempt to fully-optimize our architecture. In particular, no fine-tuning of hyper-parameters has been carried out using a validation set approach, something we leave for future work with larger datasets. Our aim has been to implement a proof of concept of the idea that deep learning approaches can provide value for the analysis of time-frequency representations of EEG data.

## 2 EEG dataset

Idiopathic RBD patients (called henceforth RBD for data analysis labeling) and healthy controls were recruited at the Center for Advanced Research in Sleep Medicine of the Hôpital du Sacrè-Cœur de Montrèal as part of another study and kindly provided for this work. All patients with a full EEG montage for resting-state EEG recording at baseline and with at least one follow-up examination after the baseline visit were included in the study. The first valid EEG for each patient enrolled in the study was considered baseline. Participants also underwent a complete neurological examination by a neurologist specialized in movement disorders and a cognitive assessment by a neuropsychologist. No controls reported abnormal motor activity during sleep or showed cognitive impairment on neuropsychological testing. The protocol was approved by the hospital’s ethics committee, and all participants gave their written informed consent to participate. For more details on the protocol and on the patient population statistics (age and gender distribution, follow up time, etc.), see [17].

As in related work [17, 19, 20], the raw data in this study consisted of resting-state EEG collected from awake patients using 14 scalp electrodes. The recording protocol consisted of conditions with periods of “eyes open" of variable duration (~ 2.5 minutes) followed by periods of “eyes closed" in which patients were not asked to perform any particular task. EEG signals were digitized with 16-bit resolution at a sampling rate of 256 S/s. The amplification device bandpass filtered the EEG data between 0.3 and 100 Hz with a notch filter at 60 Hz to minimize line power noise. All recordings were referenced to linked ears. The dataset includes a total of 121 patients diagnosed with RBD (of which 118 passed the first quality tests) and 85 healthy controls (of which only 74 provided sufficient quality data) without sleep complaints and in which RBD was excluded. EEG data was collected in every patient at baseline, i.e., when they were still RBD. After 1–10 years of clinical follow-up and, 14 patients converted to PD, 13 to DLB, while the rest remained idiopathic RBD. Our description here focuses on the previously studied balanced HC vs. PD binary classification problem.

### 2.1 Preprocessing and generation of spectrograms

To create spectrograms (the frames), EEG data from each channel was processed using Fourier analysis (FFT) after detrending blocks of 1 second with a Hann window (FFT resolution is 2 Hz) (see Figure 1). To create the spectrogram frames, 20 second 14 channel artifact-free epochs were collected for each subject, using a sliding window of 1 second. FFT amplitude bins in the band 4–44 Hz were used. The data frames are thus multidimensional arrays of the form [channels (14)] x [FFTbins (21)] x [Epochs (20)]. The number of frames per subject was fixed, as a trade-off between data per subject and number of subjects included, to 148, representing about 2.5 minutes of data. We selected a minimal suitable number of frames per subject so that each subject provided the same number of frames. For training, datasets were balanced for subjects by random replication of subjects in the class with fewer subjects. For testing, we used a leave-pair-out strategy (LPO [2]), with one subject from each class. Thus, both the training and test sets were balanced both in terms of subjects and frames per class. Finally, the data was centered and normalized to unit variance for each frequency and channel.

**Figure 1:**
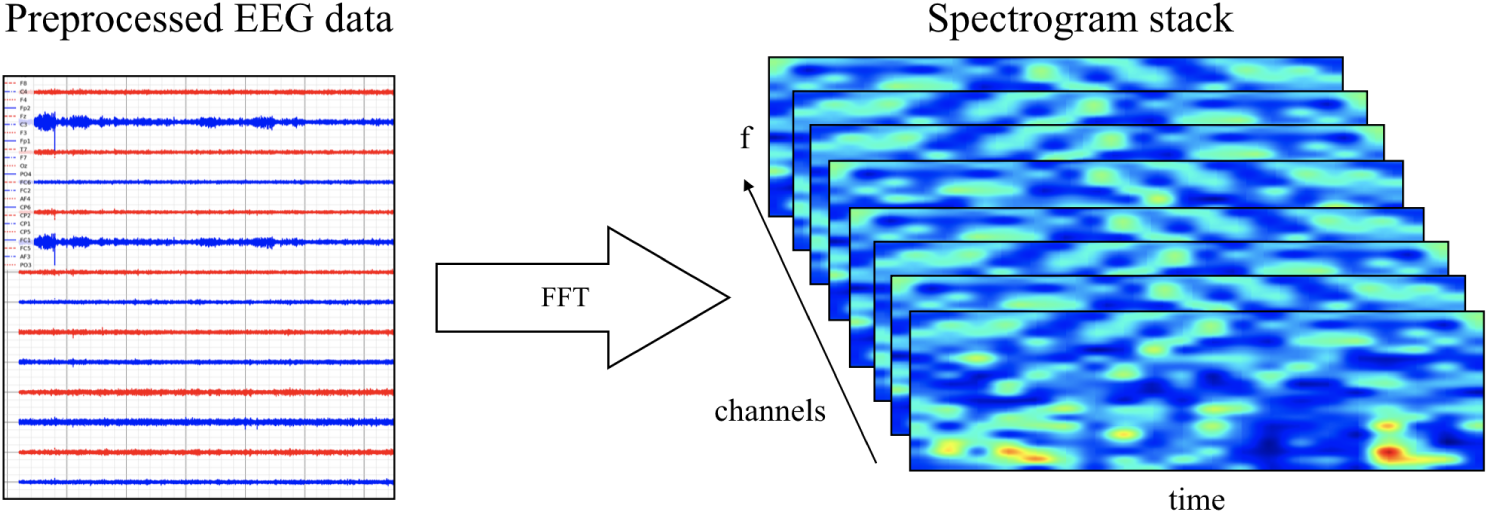
Generation of spectrogram stack for each data epoch for a subject from preprocessed (artifact rejection, referencing, detrending) EEG data.

## 3 Architectures

We have implemented three architectures: DCNN and stacked RNN, as we now describe, plus a shallow architecture for comparison—see Figure 2.

**Figure 2:**
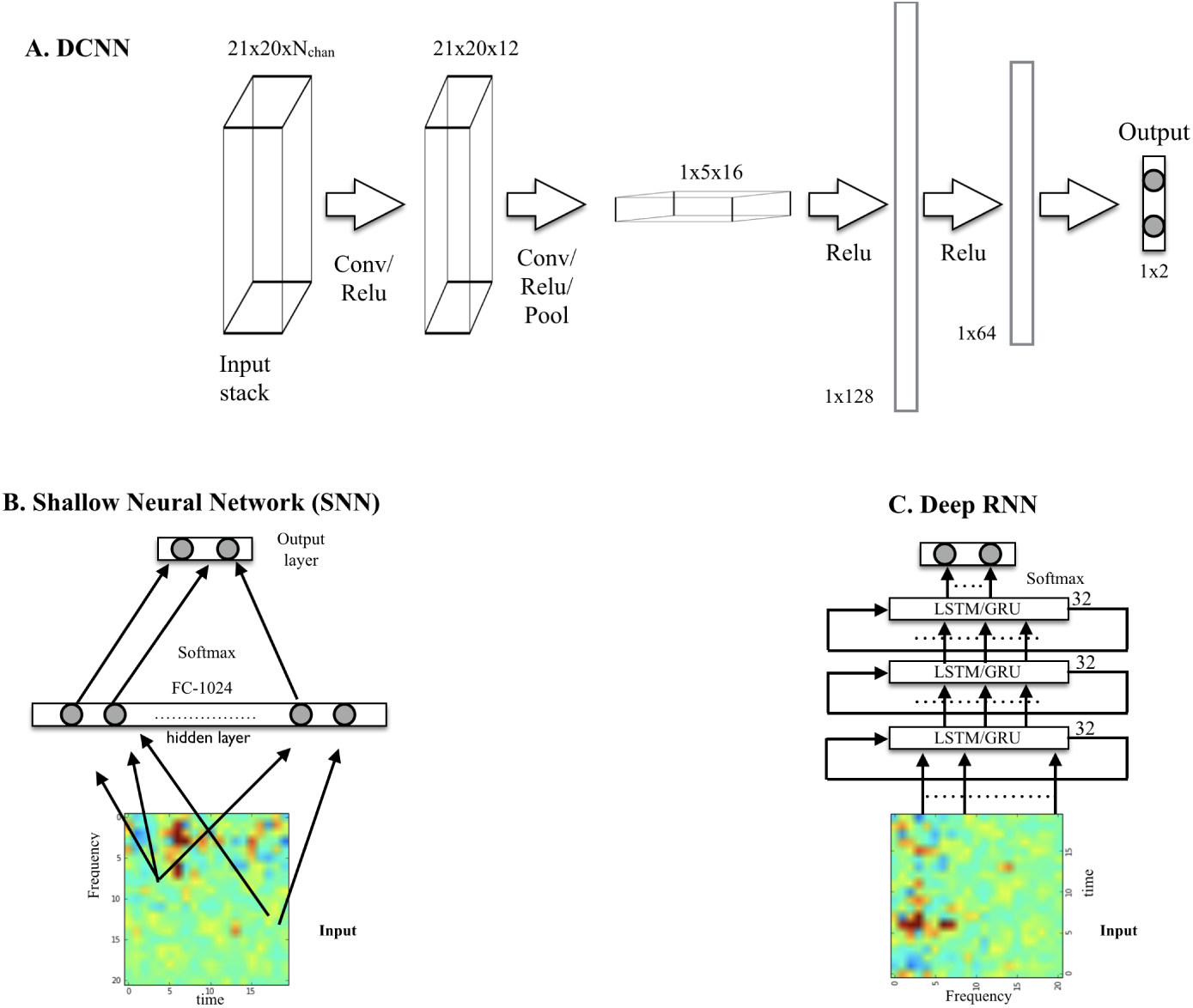
A. DCNN model displaying input, convolution with pooling layers, and hidden-unit layers. The output, here for the binary classification problem using one-hot encoding, is a two node layer. B. Shallow neural network architecture used for comparison. C. Deep RNN using LSTM or GRU cells.

### 3.1 Convolutional network architecture

The network (which we call *SpectNet)*, implemented in *Tensorflow* [1]), is a relatively simple four hidden-layer convolutional net with pooling (see Figure 2). Dropout has been used as the only regularization. All EEG channels may be used in the input cube. The design philosophy has been to enable the network to find local features first and create larger views of data with some temporal (but not frequency) shift invariance via max-pooling.

The network has been trained using a cross-entropy loss function to classify frames (not subjects). It has been evaluated both on frames and, more importantly, on subjects by averaging subject frame scores and choosing the maximal probability class, i.e., using a 50% threshold. For development purposes, we have also tested the performance of this DCNN on a synthetic dataset consisting of Gaussian radial functions randomly placed on the spectrogram time axis but with variable stability in frequency, width and amplitude (i.e, by adding some jitter top these parameters). Frame classification accuracy was high and relatively robust to jitter (~95–100%, depending on parameters).

### 3.2 RNN network architecture

The architectures for the RNNs consisted of stacked LSTM [8, 7] or GRU cells [3]. The architecture we describe here consists of three stacked cells, where each cell uses as input the outputs of the previous one. Each cell used 32 hidden units, and dropout was used to regularize it. The performance of LSTM and GRU variants was very similar.

## 4 Classification performance assessment

Classification performance has been evaluated in two ways using leave-pair out cross-validation (LPO) with a subject from each class. First, by working with balanced dataset using the accuracy metric (probability of good a classification), and second, by using the area under the curve (AUC) [2]. To map out the classification performance of the DCNN for different parameter sets, we have implemented a set of algorithms based on the *Tensorflow* package [1] as described in the following pseudocode:

~~~
REPEAT N times (experiments):
1- Choose (random, balanced) training and test subject sets (leave-pair-out)
2- Augment smaller set by random replication of subjects
3- Optimize the NN using stochastic gradient descent with frames as inputs
4- Evaluate per-frame performance on training and test set
5- Evaluate per-subject performance averaging frame outputs
END
Compute mean and standard deviation of performances over the N experiments
~~~

For each frame, the classifier outputs the probability of the frame belonging to each class (using *softmax*, see, e.g., [7]) and, as explained above, after averaging over frames per subject we obtain the probability of the subject belonging to each class. This provides an interesting score in itself. Classification is carried out by choosing the class with maximal probability.

The results from classification are shown in Table 1 for the HC vs. PD problem and the HC+RBD vs. PD+DLB problem, which includes more data. Sample results for the RNN architecture (which are very similar to DCNN results) are provided in Figure 4. For comparison, using a shallow architecture neural network resulted in about 10% less ACC or AUC (in line with our results using support vector machine (SVM) classifiers [24, 23]). Figure 5 provides the performance in the HC vs. PD problem using different EEG channels using a smaller number of folds.

**Table 1:** Performance in different problems using a single EEG channel (P4). From left to right: architecture used and problem addressed (groups); Number of subjects in training and test sets per group (always balanced); train and test average performance on frames; test accuracy and LPO cross-validation area-under-the-curve metric (AUC) [2]. Results to ±1%.

| **Problem** | **N**<br>**train/test** | **Frame**<br>**train/test**<br>**ACC** | **Subject test**<br>**ACC (AUC)** |
| --- | --- | --- | --- |
| DCNN: HC vs PD | 2x73 / 2x1 | 80% / 73% | 79% (87%) |
| RNN: HC vs PD | 2x73 / 2x1 | 77% / 74% | 81% (87%) |
| DCNN: HC+RBD vs PD+DLB | 2x159 / 2x1 | 73% / 68% | 73% (78%) |
| RNN: HC+RBD vs PD+DLB | 2x159 / 2x1 | 76% / 68% | 72% (77%) |

**Figure 3:**
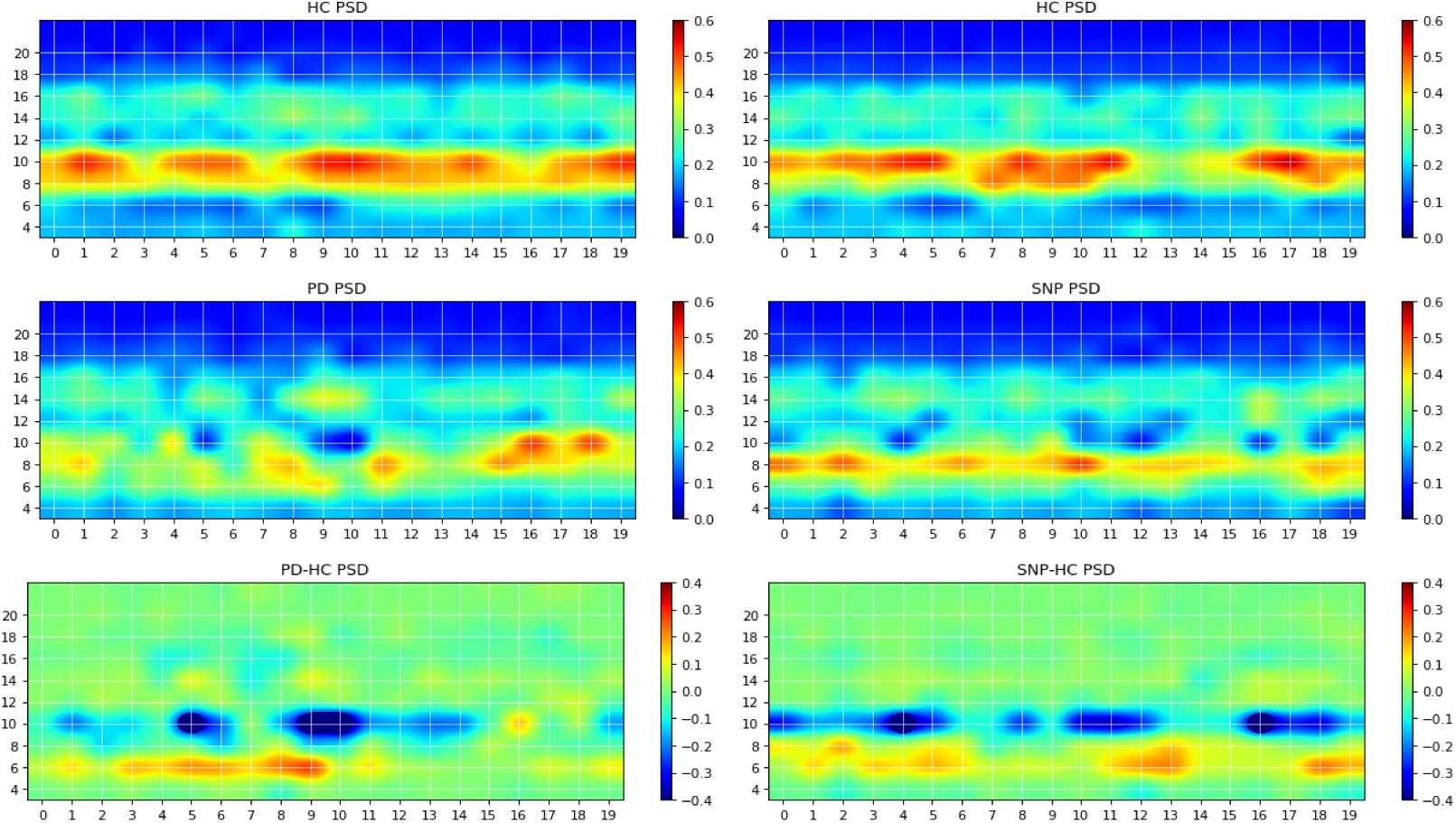
Sample images produced by maximizing network outputs for a given class. Left column: from a network was trained using P4 electrode channel data on the problem of HC vs PD. The main features are the presence of 10 Hz bursts in the HC image (top) compared to more persistent 6 Hz power in the pathological spectrogram (middle). The difference of the two is displayed at the bottom. Right column: network was trained using P4 electrode data on the problem of HC vs PD+DLB (i.e., HC vs RBDs that will develop an *α*-synucleinopathy or SNP). The main features are the presence of 10 Hz bursts in the HC image (top) compared to more persistent 6–8 Hz power in the pathological spectrogram (middle).

**Figure 4:**
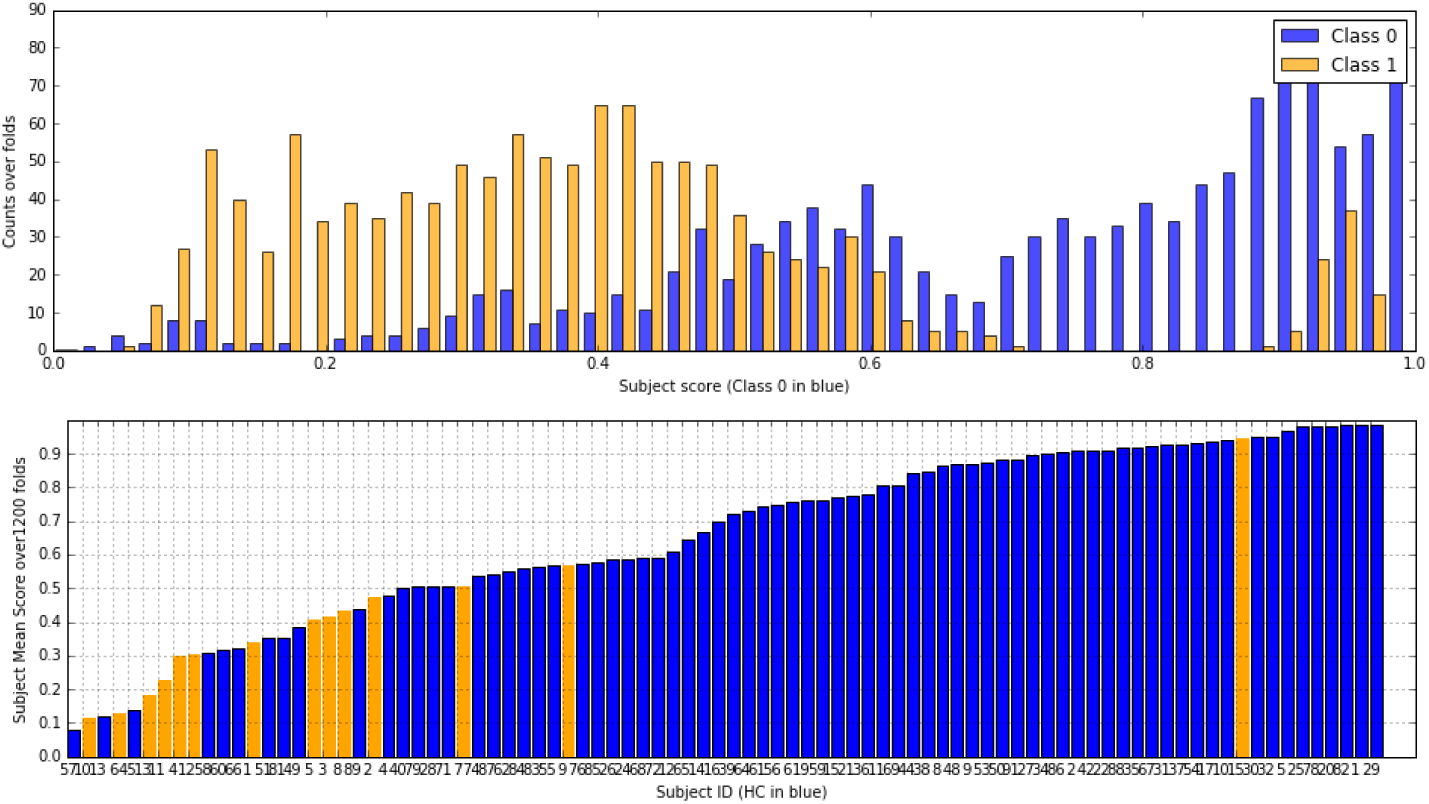
Top: RNN Frame score histogram per class (HC in blue, PD in orange) using one channel (P4). B. Subject mean score across folds. In this particular run, with mean ACC=80%, AUC=87% (both ±1%). There are clearly some subjects that are not classified correctly (this is consistently with DCNN results). The PD outlier is unusual in terms of other metrics, such as slow2fast ratio (EEG slowing) or LZW complexity [20]. Results from the DCNN are very similar.

**Figure 5:**
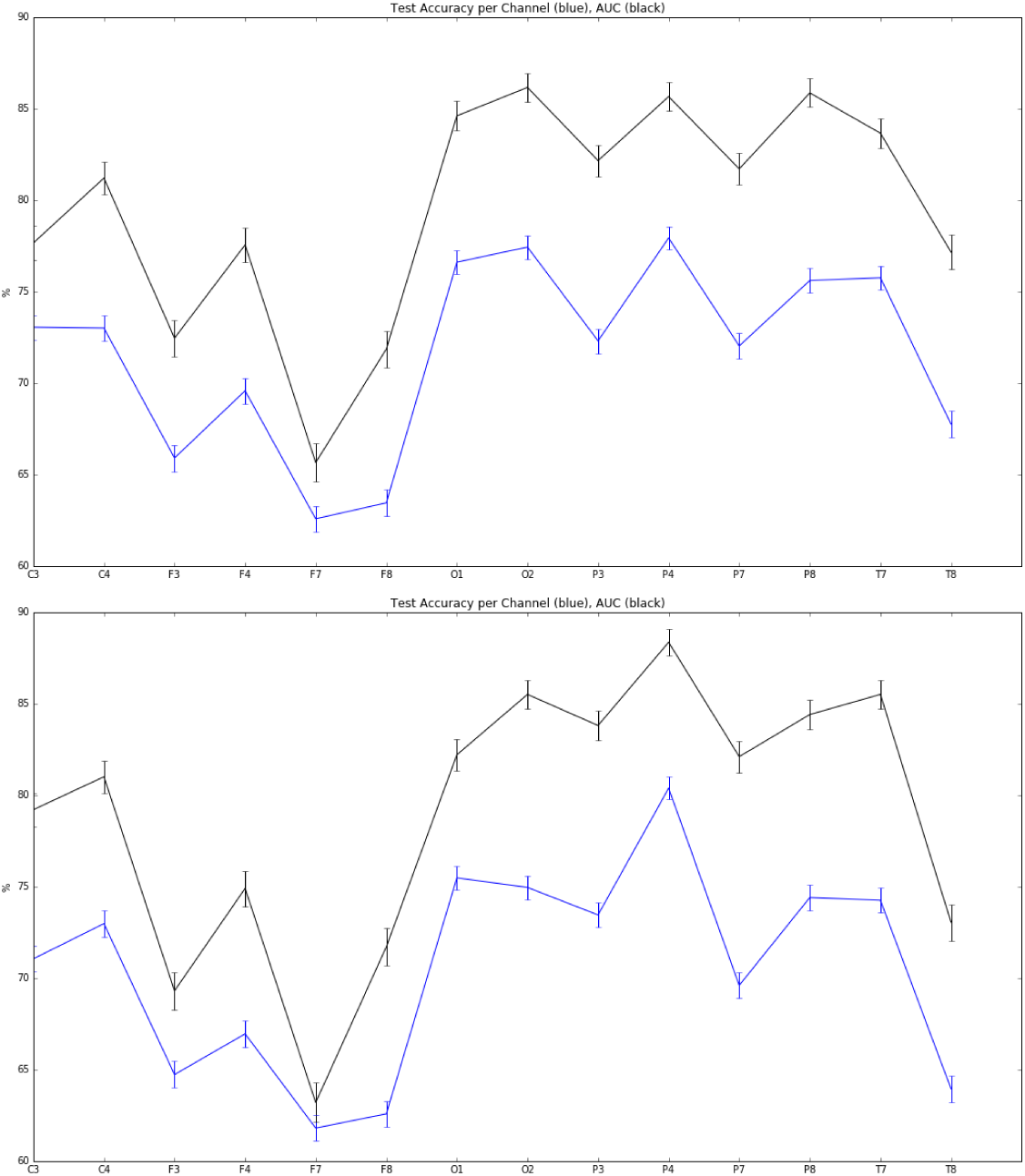
Mean accuracy and AUC per EEG channel (averages and standard error of the mean over 2000 folds) for the single channel HC vs. PD classification problem. Occipital and parietal electrodes provide better discrimination (top: DCNN architecture, bottom: RNN).

## 5 Interpretation

Once a DCNN has been trained, it can be used to explore which inputs optimally excite network nodes, including the outputs [22]. The algorithm for doing this consists essentially in maximizing a particular class score using gradient descent, starting from, e.g., a random noise image. An example of the resulting images using the DCNN above can be seen in Figure 3. This is a particularly interesting technique in our diagnosis/prognosis problem, as it provides new insights on the class-specific features in EEG of each class. In the case of a HC vs. PD trained network, we can see alterations in the alpha and theta spectral bands, appearing differentially in the form of bursts in each class.

## 6 Discussion

Our results using deep networks are complementary to earlier work with this type of data using SVMs and ESNs. However, we deem this approach to be particularly interesting for various reasons. First, it largely mitigates the need for feature selection (spectral bands and channels). Secondly, results represent an improvement over related prior efforts, increasing performance by about 5–10% in AUC [24, 23].

We note that one of the potential issues with our dataset is the presence of healthy controls without follow up, which may be a confound. We hope to remedy this by enlarging our database and by improving our diagnosis and follow up methodologies. In addition to dataset quality improvements, future steps include the exploration of this approach with larger datasets as well as a more systematic study of network architecture and regularization schemes. This includes the use of deeper architectures, improved data augmentation methods, alternative data segmentation and normalization schemes. With regard to data preprocessing, we should consider improved spectral estimation using more advanced techniques such as state-space estimation and multitapering—as in [11], and working with cortically or scalp-mapped EEG data prior creation of spectrograms.

Although here, as in [28], we worked with time-frequency pre-processed data, the field will undoubtedly steer towards working with raw data in the future when larger datasets become available—as suggested in [21]. In closing, we note that the techniques used here can be extended to other EEG related problems, such as brain-computer interfaces, sleep scoring, detection of epileptiform activity or EEG data pre-processing, where the advantages of deep learning approaches may prove useful as well.

## References

[1] Martin Abadi, Paul Barham, Jianmin Chen, Zhifeng Chen, Andy Davis, Jeffrey Dean, Matthieu Devin, Sanjay Ghemawat, Geoffrey Irving, Michael Isard, Manjunath Kudlur, Josh Levenberg, Rajat Monga, Sherry Moore, Derek G. Murray, Benoit Steiner, Paul Tucker, Vijay Vasudevan, Pete Warden, Martin Wicke, Yuan Yu, and Xiaoqiang Zheng. Tensorflow: A system for large-scale machine learning. In 12th USENIX Symposium on Operating Systems Design and Implementation (OSDI 16), pages 265–283, 2016.

[2] A. Airola, T. Pahikkala, W. Waegeman, B. De Baets, and T. Salakoski. A comparison of AUC estimators in small-sample studies. In Journal of Machine Learning Research - Proceedings Track, pages 3–13, 2010.

[3] Kyunghyun Cho, Bart van Merrienboer, Caglar Gulcehre, Fethi Bougares, Holger Schwenk, and Yoshua Bengio. Learning phrase representations using RNN encoder-decoder for statistical machine translation. In Conference on Empirical Methods in Natural Language Processing (EMNLP 2014), arXiv preprint arXiv:1406.1078, 2014.

[4] J. Dauwels, F. Vialatte, T. Musha, and A. Cichocki. A comparative study of synchrony measures for the early diagnosis of Alzheimer’s disease based on EEG. NeuroImage, (49):668–693, 2010.

[5] L. Fantini, J.F Gagnon, D. Petit, S. Rompre, A. Decary, J. Carrier, and J. Montplaisir. Slowing of electroencephalogram in rapid eye movement sleep behavior disorder. Annals of neurology, 53(6):774–780, 2003.

[6] Stephany Fulda. Idiopathic REM sleep behavior disorder as a long-term predictor of neurodegenerative disorders. EPMA J., 2(4):451–458, December 2011.

[7] Ian Goodfellow, Yoshua Bengio, and Aaron Courville. Deep Learning. MIT Press, 2016.

[8] Sepp Hochreiter and Jürgen Schmidhuber. Long short-term memory. Neural Computation, 9(8):1735–1780, Nov 1997.

[9] Birgit Högl, Ambra Stefani, and Aleksandar Videnovic. Idiopathic REM sleep behavior disorder and neurodegeneration — and update. Nature Review Neurology, 14(1):40–55, 2018.

[10] Alex Iranzo, Ana Fernández-Arcos, Eduard Tolosa, Mónica Serradell, José Luis Molinuevo, Francesc Valldeoriola, Ellen Gelpi, Isabel Vilaseca, Raquel Sánchez-Valle, Albert Lladó, Carles Gaig, and Joan Santamaría. Neurodegenerative disorder risk in idiopathic REM sleep behavior disorder: Study in 174 patients. PLoS One, 9(2), February 2014.

[11] Seong-Eun Kim, Michael K. Behr, Demba Ba, and Emery N. Brown. State-space multitaper time-frequency analysis. PNAS, www.pnas.org/cgi/doi/10.1073/pnas.1702877115(1702877115), 2017.

[12] Yoon Kim, Young Eun Kim, Eun Ok Park, Chae Won Shin, Han-Joon Kim, and Beomseok Jeon. REM sleep behavior disorder portends poor prognosis in Parkinson’s disease: A systematic review. J Clin Neurosci., Oct 2017.

[13] E Kroupi, A Soria-Frisch, M Castellano, D Ibá nez, J Montplaisir, J-F Gagnon, R Postuma, S Dunne, and G Ruffini. Deep networks using auto-encoders for PD prodromal analysis. In Proceedings of 1st HBP Student Conference, Vienna, 2017.

[14] R. Laje and D.V. (2013). D.V. Buonomano. Robust timing and motor patterns by taming chaos in recurrent neural networks. Nat Neurosci, 16:925–933, 2013.

[15] Tomaso Poggio, Hrushikesh Mhaskar, Lorenzo Rosasco, Brando Miranda, and Qianli Liao. Why and when can deep but not shallow networks avoid the curse of dimensionality: a review. Technical Report CBMM Memo 058, Center for Brains, minds and machines, 2016.

[16] RB Postuma, JF Gagnon, M Vendette, ML Fantini, J Massicotte-Marquez, and et al. Quantifying the risk of neurodegenerative disease in idiopathic REM sleep behavior disorder. Neurology, 72:1296–1300, 2009.

[17] J Rodrigues-Brazète, JF Gagnon, RB Postuma, JA Bertrand, and Montplaisir J D Petit. Electroencephalogram slowing predicts neurodegeneration in rapid eye movement sleep behavior disorder. Neurobiol Aging, 37(74–81), Jan 2016.

[18] G Ruffini. Models, networks and algorithmic complexity. Starlab Technical Note - arXiv:1612.05627, TN00339(DOI: 10.13140/RG.2.2.19510.50249), December 2016.

[19] Giulio Ruffini, David Ibañez, Marta Castellano, Stephen Dunne, and Aureli Soria-Frisch. EEG-driven RNN classification for prognosis of neurodegeneration in at-risk patients. ICANN 2016, 2016.

[20] Giulio Ruffini, David Ibañez, Eleni Kroupi, Jean-François Gagnon, Jacques Montplaisir, Ronald B. Postuma, Marta Castellano, and Aureli Soria-Frisch. Algorithmic complexity of EEG for prognosis of neurodegeneration in idiopathic rapid eye movement behavior disorder (RBD). bioRxiv, (200543), 2017.

[21] Robin Tibor Schirrmeister, Jost Tobias Springenberg, Lukas Dominique Josef Fiederer, Martin Glasstetter, Katharina Eggensperger, Michael Tangermann, Frank Hutter, Wolfram Burgard, and Tonio Ball. Deep learning with convolutional neuralnetworks for eeg decoding and visualization. Human Brain Mapping, 38:5391–5420, 2017.

[22] Karen Simonyan, Andrea Vedaldi, and Andrew Zisserman. Deep inside convolutional networks: Visualising image classification models and saliency maps. arXiv:1312.6034 [cs.CV], 2014.

[23] A. Soria-Frisch, J. Marin, D. Ibañez, S. Dunne, C. Grau, G. Ruffini, J. Rodrigues-Brazète, R. Postuma, J.-F. Gagnon, J. Montplaisir, and A. Pascual-Leone. Evaluation of EEG biomarkers for Parkinson disease and other Lewy Body diseases based on the use of machine learning techniques. in preparation, 2018.

[24] A. Soria-Frisch, J. Marin, D. I Ibañez, S. Dunne, C. Grau, G. Ruffini, J. Rodrigues-Brazète, R. Postuma, J.-F. Gagnon, J. Montplaisir, and A. Pascual-Leone. Machine learning for a Parkinson’s prognosis and diagnosis system based on EEG. Proc. International Pharmaco-EEG Society Meeting PEG 2014, Leipzig, Germany, 2014.

[25] Orestis Tsinalis, Paul M. Matthews, Yike Guo, and Stefanos Zafeiriou. Automatic sleep stage scoring with single-channel EEG using convolutional neural networks. arXiv:1610.01683v1, 2016.

[26] Peter J. Uhlhaas and Wolf Singer. Neural synchrony in brain review disorders: Relevance for cognitive dysfunctions and pathophysiology. Neuron, 52:155–168, 2006.

[27] A. van den Oord, S. Dieleman, and Benjamin Schrauwen. Deep content-based music recommendation. In *NIPS*, 2013.

[28] Albert Vilamala, Kristoffer H. Madsen, and Lars K. Hansen. Deep convolutional neural networks for interpretable analysis of EEG sleep stage scoring. In 2017 IEEE International Workshop On Machine Learning For Signal Processing, Sept. 25–28, 2017, Tokyo, Japan, 2017.

